# Repetitive DNA content in the maize genome is uncoupled from population stratification at SNP loci

**DOI:** 10.1101/652859

**Authors:** Simon Renny-Byfield, Andy Baumgarten

## Abstract

**Motivation:** Repetitive DNA is a major component of plant genomes and is thought to be a driver of evolutionary novelty. Describing variation in repeat content among individuals and between populations is key to elucidating the evolutionary significance of repetitive DNA. However, the cost of producing references genomes has limited large-scale intraspecific comparisons to a handful of model organisms where multiple reference genomes are available.

**Results:** We examine repeat content variation in the genomes of 94 elite inbred maize lines using graph-based repeat clustering, a reference-free and rapid assay of repeat content. We examine population structure using genome-wide repeat profiles and demonstrate the stiff-stalk and non-stiff-stalk heterotic populations are homogenous with regard to global repeat content. In contrast and similar to previously reported results, the same individuals show clear differentiation, and aggregate into two populations, when examining population structure using genome-wide SNPs. Additionally, we develop a novel kmer based technique to examine the chromosomal distribution of repeat clusters *in silico* and show a cluster dependent statistically significant association with gene density.

**Conclusion:** Our results indicate that repeat content variation in the heterotic populations of maize has not diverged and is uncoupled from population stratification at SNP loci. We also show that repeat families exhibit divergent patterns with regard to chromosomal distribution, some repeat clusters accumulate in regions of high gene density, whereas others aggregate in regions of low gene density.

**Author’s contributions:** SRB and AB conceived the study, SRB performed the bioinformatic analysis, SRB wrote the paper with input from AB. email contacts: Simon Renny-Byfield, Andy Baumgarten

## 1 Introduction

Maize is a commercially important hybrid crop grown globally for livestock consumption, food, and fuel. Early breeding programs observed that crosses between maize populations resulted in increased heterosis where a F1 progeny or hybrid demonstrates significantly higher performance than its parents [1, 2]. Public and private industry breeding programs have exploited heterosis in maize to maximize genetic gain for grain yield [3]. These programs employ reciprocal recurrent selection to develop specific germplasm pools, called heterotic groups, that maximize heterosis in maize. Improvement occurs within heterotic groups to develop inbred lines that maximize heterosis in crosses between heterotic groups. These inbred lines are then recycled to create new crosses within each heterotic pool.

Several studies have demonstrated that reciprocal recurrent selection drives differences in allele frequencies between heterotic groups. These studies have coupled genotyping and population genetics approaches to demonstrate there are three distinct heterotic pools within current North American elite maize germplasm. These pools, referred to as Stiff-Stalk (SS), Non-Stiff-Stalk (NSS), and Iodent (IOD) show increasing allele frequency divergence over time with recently developed lines having the greatest divergence.

Previous studies examining the genetic divergence, allelic frequency, and linkage disequilibria between maize heterotic pools have focused solely on SNPs found in genic regions of the maize genome. However, the maize genome consists primarily of various repetitive DNA families, similar to other species with similar genome size. Studies have demonstrated that repetitive DNA dynamics can influence gene content changes [4], changes in genome size [5], and gene expression activation within plants. [6, 7] A study comparing the divergence and abundance of repeat variation between maize heterotic pools could provide insight to whether repeats follow the same divergence as SNP-based diversity studies.

Unfortunately, the cost and complexity of producing high-quality reference genomes limits reference-based comparisons of repeat content to a few species and individuals with fully sequenced genomes. These cost and analysis restrictions limit the ability to compare repeat content and its association to population and phenotypic variation. However, reference-free approaches using low coverage short-read sequencing and graph-based repeat clustering have been used to efficiently assay repeat content in eukaryotic genomes[8]. Reference-free, repeat-based methods can exploit this repeat sequence abundance to estimate repeat sequence similarity and abundance across hundreds of different genomes without the use of a genomic reference. [9, 10, 11, 12, 13]. The highly repetitive nature of the maize and other plant genomes ensures that the majority of genomic reads coming from short-read sequencing will be repetitive in nature.

Reference-free approaches, using skim sequencing and repeat clustering, were successfully used to assaying repeat content in allopolyploid tobacco and demonstrated the biased removal of repetitive DNA from one of the two sub-genomes[12]. Furthermore, similar clustering, along with empirical validation, was used to demonstrate the rapid elimination of tandem repeat sequences in synthetic allopolyploids[9]. Other studies in *Nicotiana* have shown reductions in genome size occurs due to the loss of low copy-number repeats while genome expansion resulted from the increase of highly abundant repeats [5]. Additionally, repeat clustering has revealed the repeat content of rye B-chromosomes [14] and the giant genomes of *Fritilaria* [10]. More importantly, repeat profiles generated by repeat-clustering exhibit phylogenetic signal consistent with that observed for plastid and nuclear markers in a variety of taxa [15].

In this study, we take advantage of skim sequencing and graph-based repeat clustering to screen the genomes of 94 elite ex-PVP maize inbreds. Ex-PVP lines are inbreds developed and patented by private or public institutions but have had their patents expire allowing public breeding and genotyping use. Many of the lines included in this study represent the base germplasm used to establish current industry and public North American maize breeding programs [16]. We identify and annotate repeat families *de novo* and examine their differential abundance within and between heterotic groups. The chromosomal distribution of repeat families was also compared between three high-quality reference maize genome (citations) using a novel k-mer based method.

Our analysis revealed no significant differentiation in abundance of the two major classes of repeats present in the maize genome, *Gypsy* and *Copia* LTR-retroelements, when comparing the two maize heterotic populations. We observed that the chromosomal distribution varies between repeat families and that statistically significant association exists between the chromosome distribution of repeat clusters and gene density. We found that SNP data from primarily genic regions shows significant population structure between the two maize heterotic pools. In contrast, similar population structure was not found using repeat clustering data, suggesting that divergence of the heterotic groups has occurred at the SNP level and that this process is not mirrored at the level of repeat abundance or genomic distribution.

## 2. Methods

### 2.1 Data Preparation and Read Sampling

Ninety four ex-PVP lines were selected for short-read sequencing. Inbred lines were categorized into stiff-stalk (SS) and non-stiff-stalk (NSS + IO) heterotic groups using previous publications or internal knowledge (citation). DNA was extracted with V5 stage leaf tissue from 94 greenhouse grown accession (see Supp Info File 1) using the CTAB method. Illumina HiSeq 2000 libraries where prepared from purified DNA according to the manufacturers instructions and samples were sequenced at 20-60x coverage using the Illumina HiSeq 2000 instrumentation resulting in 150 bp paired-end reads. For each sample, the resulting sequencing data were randomly down-sampled, resulting in a skim sequence dataset of 50,000 whole genome shotgun (WGS) reads per individual, taking only one read from a pair. Sequence reads are available at the NCBI SRA under the BioProject number PRJNA530574.

### 2.2 Graph-based Repeat Clustering

We implemented graph-based clustering, based on previously published methods[8], using data from all sequenced ex-PVP inbreds as input. The method from [8] was re-written in Python 2.7 using the Python version of igraph[17] to increase analysis speed. Relevant code is available on request. Graph-based repeat clusters was performed by pooling sequencing reads from all ex-PVP inbreds into a single dataset. Sequence reads from each individual were tracked in the combined pool to allow each read to be linked to the individual inbred it came from. Repeat families were then identified using a graph-based clustering approach similar to previously described methods [8].

Briefly, a complete pair-wise comparison is performed between all reads using megablast [18]. Using this data a simple, undirected graph of the form

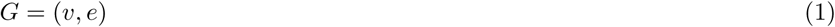

was constructed from the resulting pair-wise relationships such that vertices (*v*) of the graph (*G*) represent reads from our dataset and edges (*e*) between the vertices correspond to megablast hits between the reads. Edges were weighted according to the bit-score of the corresponding blast hit and edges with bit-score less than 100 were removed from further analysis. The graph object was reduced to its largest connected component and a community detection process was performed using the fast greedy algorithm [19] to detect repeat families. This process of community detection established clusters of reads closely connected in the graph representing highly-related repetitive DNA families. The number of reads contributed from the sequence of each ex-PVP inbred was quantified for each cluster. For the 25 largest clusters we extracted the cluster sub-graph *G*_*s*_ from the graph *G* and derived the sub-graph layout using the Fruchterman and Reingold algorithm [20], positioning vertices (reads) such that reads containing similar sequence are placed close together in 2D space.

We generated a cluster graph (*G*_*c*_) from the graph *G* of the form:

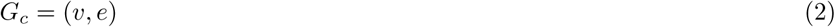

where vertex *v*_*i*_ represents the *i* th cluster, and the edge *e*(*v*_*i*_, *v*_*j*_) represent the sum edge weight of all edges between *v*_*i*_ and *v*_*j*_ in the graph *G*. This graph describes the relationships and connections between clusters, where each cluster has been reduced to a single vertex.

### 2.3 Repeat Cluster Annotation

Each cluster was annotated by extracting all sequence reads and using Repeat-Masker [21], with the cross match option, to search against the *plant collections* of the RepBase database [22]. Each cluster was then defined by the most common repeat match. We counted all the reads attributed to each cluster, and each annotation, on a per individual basis. We examined for statistical differences in mean cluster and mean annotation abundance in the SS and NSS population using a two-sided t-test, correcting for multiple comparisons using the method of Benjamini and Hockberg [23].

### 2.4 K-mer Analysis and Chromosome Painting

For each of the 25 largest clusters we identified all the k-mers present in the reads from that cluster using JellyFish [24] and identified the 100 most abundant k-mers present in each individual cluster. From each of te largest 25 clusters the 100 most abundant k-mers were mapped to the B73 RefGen_v4, B73 RefGen_v3.23 and the PH207 reference genomes (all downloaded from https://www.maizegdb.org/ accessed 19/9/2017) using the *fuzznuc* function of the EMBOSS package [25]. We then counted the number of k-mers mapping to 200 kb windows of the genome (step-size 100 kb) using bedtools, scaled the data and loess smoothed over profiles using the scipy package in python 2.7. For each reference sequence we generated gene density traces using the *coverage* tool of the *bedtools* package over the same 200 kb windows of the genome. Using the scaled data for each of the 25 largest clusters, we calculated the Pearson’s and Spearman’s correlation coefficient between gene density and kmer mapping density using the scipy package in python 2.7.

### 2.5 PCA Analysis

Using cluster abundance estimates from each of the 94 inbreds, we performed PCA analysis. Abundance measures were normalized on a cluster by cluster basis using minimum/maximum scaling of the statistics package sklearn in python 2.7, such that values for each cluster ranged from zero to one. Using the normalized data we fit a PCA with 2 components using functionality of the sklearn package in Python 2.7. We performed the same analysis, but in place of cluster abundances used 1585 genome-wide SNP markers, encoding the major allele as 1, the minor allele as 0, and masking heterozygous sites.

## 3 Results and Discussion

### 3.1 Graph-Based Repeat Clustering

Graph-based repeat clustering is a computational approach that identifies and quantifies repeat families within a genome without the need for a reference genome. The approach utilizes low-coverage sequence data to construct a graph of relationships between reads based on sequence similarity. This graph was analyzed using community detection approaches to group reads into clusters that represent different repeat families. These methods were used to analyze low coverage sequence from 94 elite, ex-PVP maize inbreds. The resulting graph (*G*) contained 2,956,096 vertices (reads) and 244,031,339 edges (sequence similarity hits between reads), with a mean degree of 165.10. Community detection using the fast greedy algorithm grouped vertices into 728 repeat clusters with the largest cluster containing 207,094 vertices and the smallest containing two.

Annotation of the clusters using the Repbase repeat library revealed a diverse collection of repeat families with the majority of repeats families categorized as either *Gypsy* or *Copia* LTR retroelements (See Supp Info File 2 and 3). These results are similar and expected given previous studies using reference quality genome assemblies in maize [26, 27]. A higher abundance of *Gypsy*-like elements was observed, when compared with *Copia*-like elements, suggesting that Gypsy-like elements are typically more abundant in the genomes of elite inbred maize lines. This was true for both the overall dataset (NSS and SS populations combined; t-statistic = 21.23, p-value < 0.0001), and when considering SS (t-statistic = 14.14, p-value < 0.0001) and NSS (t-statistic = 15.83, p-value ¡ 0.0001) populations separately. No significant difference was seen between the abundance of *Gypsy*-like sequences between SS and NSS inbreds (t-statistic = 15.83, p-value= 0.38). However, marginally significant differences were observed in the total abundance of *Copia*-like repeats between SS and NSS inbreds (t-statistic = −2.12, p-value = 0.045).

We tested for significant differences in the abundance of individual repeat clusters between the NSS and SS populations. After correcting for multiple comparisons, we found that only three of the 728 clusters exhibited evidence of differential abundance between SS and NSS inbreds. This observation suggests that the NSS and SS populations are very similar in terms of global repeat abundance profiles. It should be stressed that the similarities in repeat abundance estimated by graph-based clustering are only representative of global repeat content in the genome. In contrast, local repeat and TE variation in and around genes is incredibly diverse between maize inbreds [28]. Such micro-scale variation is not reflected in our global repeat profiles.

The observation of little to no differentiation between maize heterotic groups is in contrast to other comparisons using the same methodology. For example, graph-based repeat clustering is able to detect changes in repeat abundance between closely related species in a number of plant families [5, 13, 11]. Other studies have identified and confirmed differential repeat content in rye accessions segregating for B-chromosomes [14]. Similarly, clustering analysis and empirical validation tracked the fate of individual repeat families in early generation synthetic allopolyploid tobacco and demonstrated alteration in the chromosomal distribution and abundance of those clusters [9]. As such, we know graph-based clustering is a suitable method to quickly analyze repeat content data, and detect differences between individuals of the same species or closely related taxa.

### 3.2 Chromosomal Distribution and Correlation with Gene Density

For each of the 25 most abundant clusters we calculated the cluster layout and placed vertices (reads) in 2D space. This analysis revealed incredible diversity in cluster layout (Fig. 2 A-D), between repeat families within the maize genome. Tightly compressed clusters represent collections of highly similar reads since each vertex is positioned based on the edge weight of connections to other vertices. This indicates uniformity in repeat sequence across the repeat cluster and may indicate a recent copy-number expansion. In contrast, clusters with a more dispersed layout contain more divergent reads and likely represent older repeat expansion. Several clusters, including cluster 17 (Fig. 2 D), exhibit a ring like network structure, a lay-out resulting in repeating units arranged in tandem, a hallmark of tandem repeat sequences. Importantly, such clusters typically exhibit highly localized chromosomal distributions, as might be expected of a tandem repeat sequence (Fig. 2 D, bottom panel).

**Figure 1:**
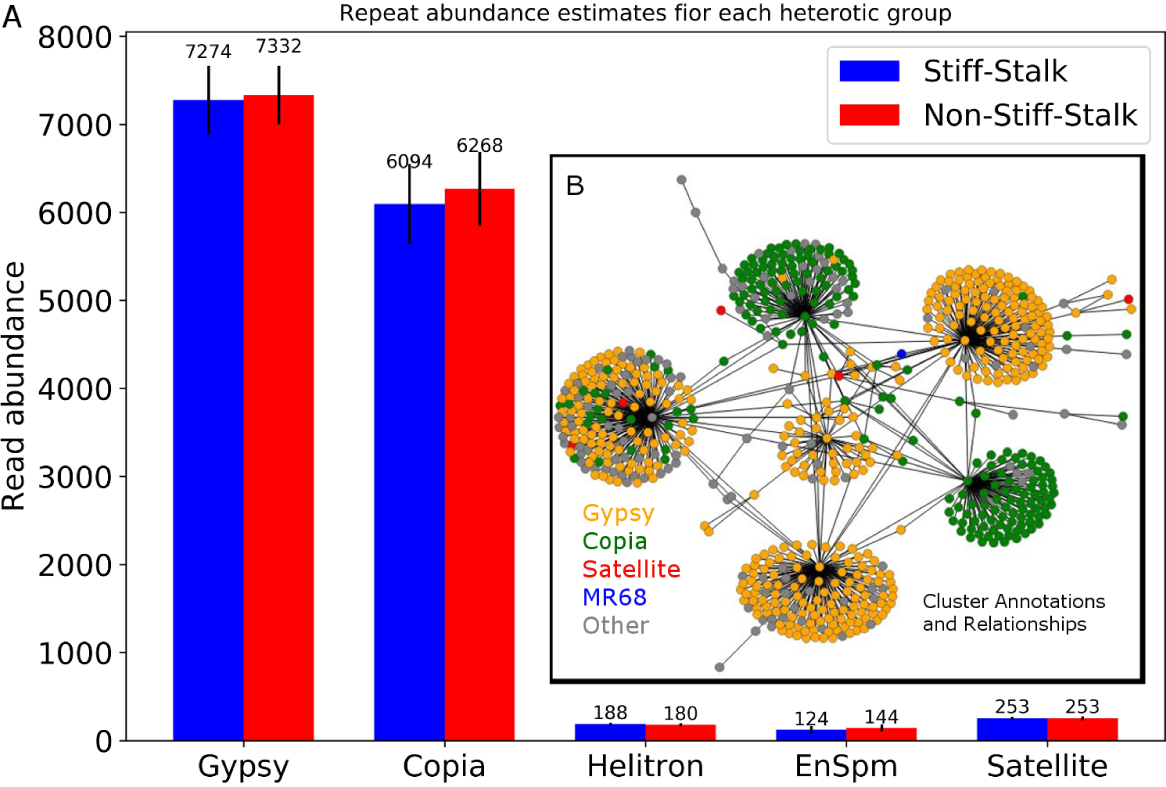
Repeat family abundance estimates and cluster relationships. (A) Using the annotation associated with each cluster we summed the total number of reads from each inbred attributed to a given annotation type, and display the mean (bars) and standard deviation (whiskers) for the SS and NSS populations for select repeat annotations. In (B) is displayed the cluster graph *G*_*c*_, where nodes are clusters and the edges indicate cross-cluster megablast hits, nodes are coloured according to annotation type.

**Figure 2:**
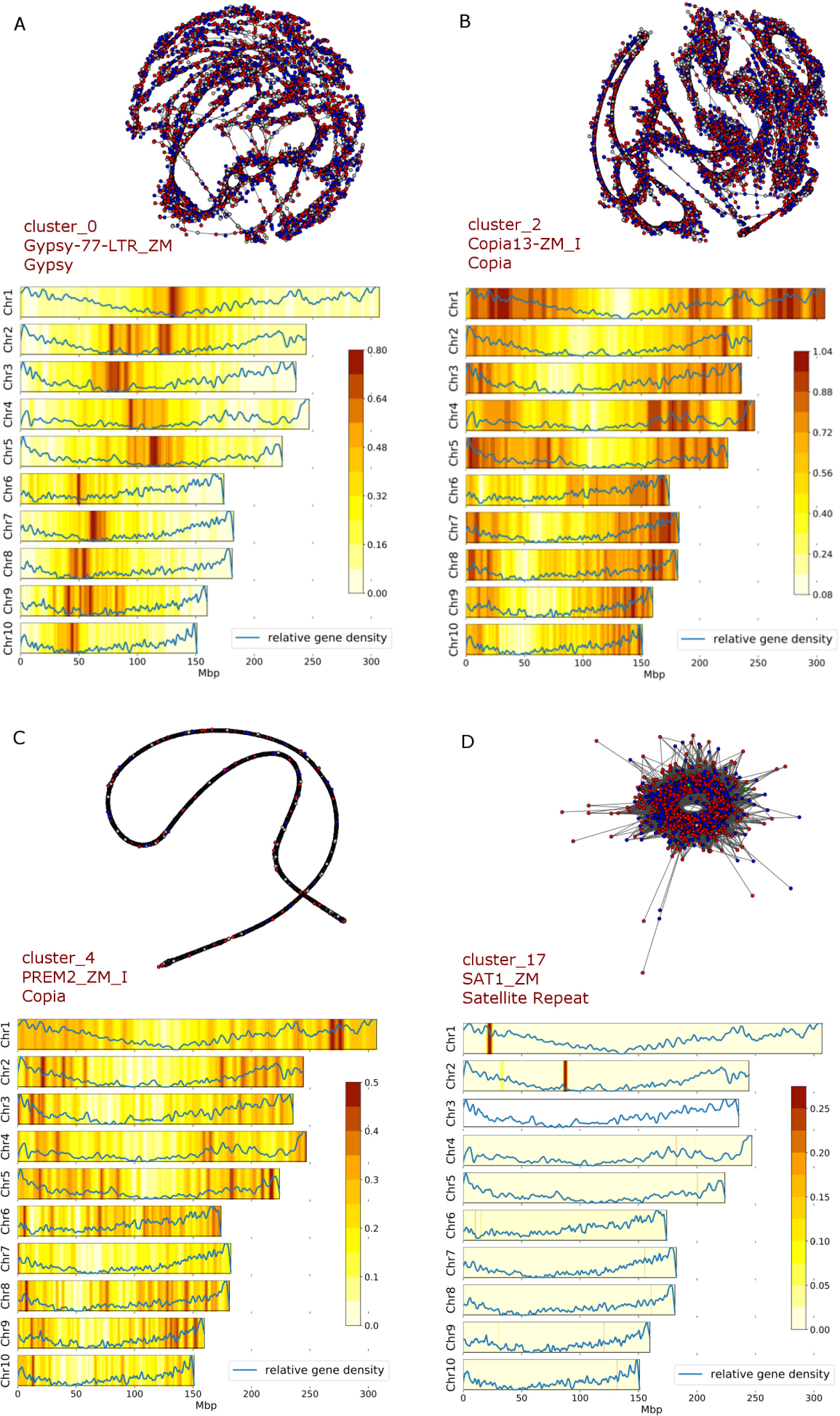
Repeat Cluster layouts and Chromosomal Distributions. Cluster layouts, in 2D space, are calculated using the Fruchterman-Reingold algorithm and are shown in the top of each panel. Reads deriving from Stiff-stalk (SS) and Non-stiff-stalk (NSS) inbreds are indicated by blue and red vertices respectively. Edges, indicated in gray, represent sequence similarity hits between reads (vertices). The cluster name, most common hit to RepBase and the repeat family to which each cluster belongs is indicated. The most abundant k-mers (of length 12, see methods) from each repeat cluster were mapped to the maize B73 RefGen_v3 genome sequence, and the resulting loess smoothed profiles for each chromosome are displayed in the lower portion of each panel. Gene density profiles are also shown in blue, overlaying the repeat profiles.

We deploy a novel *in silico* chromosome painting technique to investigate the chromosomal distribution of individual clusters. We identified the 100 most common k-mer sequences within each cluster and mapped these to the genome (Fig. 2 and Supp Info File 4) to establish the chromosomal distribution of each repeat cluster. Repeats from several clusters preferentially mapped to gene poor pericentromeric regions (Fig. 2 A) whereas k-mers from other clusters map to gene rich regions of the genome, and have lower abundance in gene poor centromeric regions (Fig. 2 B). We quantified this trend by determining the correlation between kmer mapping density and regions of the genome identified as genic (Fig. 3). 44 of the 50 largest repeat clusters showed significantly significant (p-value < 0.05) association with gene density using both methods used in this study. Although most clusters have statistically significant association with gene density we note that the coefficient is small and statistical significance is achieved through large N. It also worth noting that while there is a positive relationship between gene density and kmer mapping density, all clusters tend not to accumulate in the most gene dense regions of the genome (Fig. 3). Using cluster 0 as an example the density of mapped kmers is significantly negatively correlated with gene density, suggesting this repeat family preferentially insert into gene poor regions of the genome, or alternatively that copies are removed more easily from genic regions (Fig. 2 and Fig. 3). Interestingly we see the opposite pattern for kmers derived from cluster 2, where a positive association is observed between kmer mapping density and gene density suggesting this repeat family aggregates in gene rich regions.

**Figure 3:**
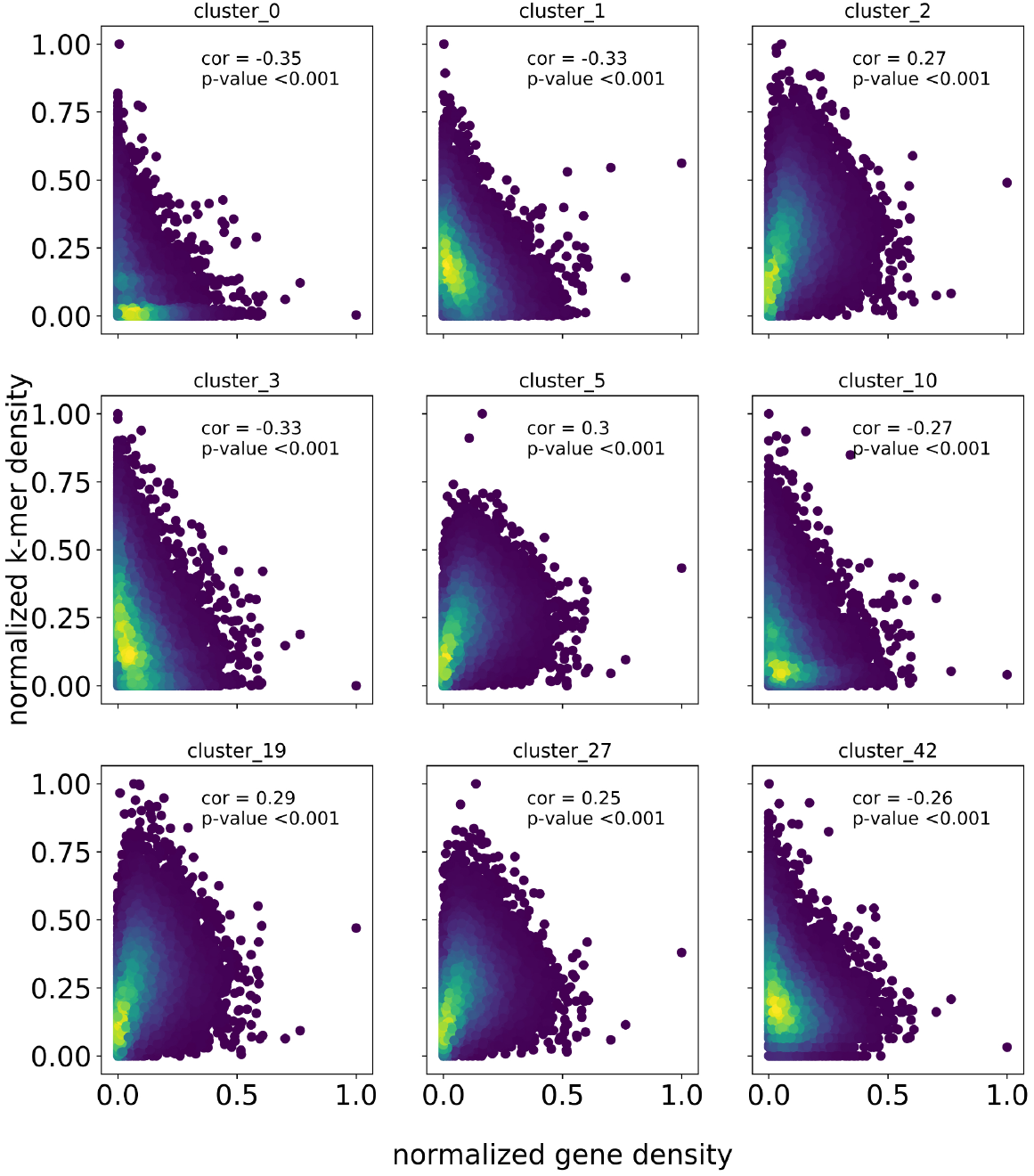
Scatter plots showing the relationship between gene density and kmer mapping density in 200 kb windows of the genome for examplar repeat clusters. For each example the Pearson’s correlation coefficient and associated p-value are given. For cluster 0, 1, 3, 10 and 42 there is a significant negative correlation, whereas a positive correlation is observed for clusters 2, 5, 19 and 27. Each data point is colored by a two dimensional kernel density estimation, where the purple to yellow color indicates ever greater density of data points.

The observation that chromosomal distribution varies between clusters and is associated to gene density is similar to previous studies on repeat sequences in maize [29] and other plant species [30, 31, 32]. For example *Ds* elements have accumulated in the sub-telomeric regions of maize chromosomes, but are relativity rare in pericentromeric regions [29]. Gypsy, CACTA, and Copia elements have varying chromosomal distribution in the *Setaria* genome [33]. More broadly, these observation likely reflect differences in the insertion preferences between repeat families and bias in the removal of repeat copies related to gene density.

### 3.3 Population Structure: A Tale of Two Datasets

We examined population structure among 94 ex-PVP inbred maize lines using global repeat abundance profiles and using standard SNP variation (Fig. 4). A PCA analysis using repeat abundance estimates revealed that ex-PVP inbreds do not appear to cluster into known heterotic groups (Fig. 4 A). The SS and NSS individuals form a single homogeneous group with SS and NSS individuals intermixed. This suggests that the SS and NSS populations are not distinct given patterns of global repeat abundance, even though repeat differences exist between individuals. In contrast, a similar PCA analysis performed using SNP information from the same inbreds separated known SS and NSS inbreds into distinguishable, significant clusters, especially on the first principle component (Fig. 4 B). The results from this analysis of SNPs are highly similar to what has been observed in previous studies using SNP data on the same and additional maize breeding and historical lines [34, 35]. The results in this study demonstrate that the population structure observed using SNPs is not reflected in global repeat abundance.

**Figure 4:**
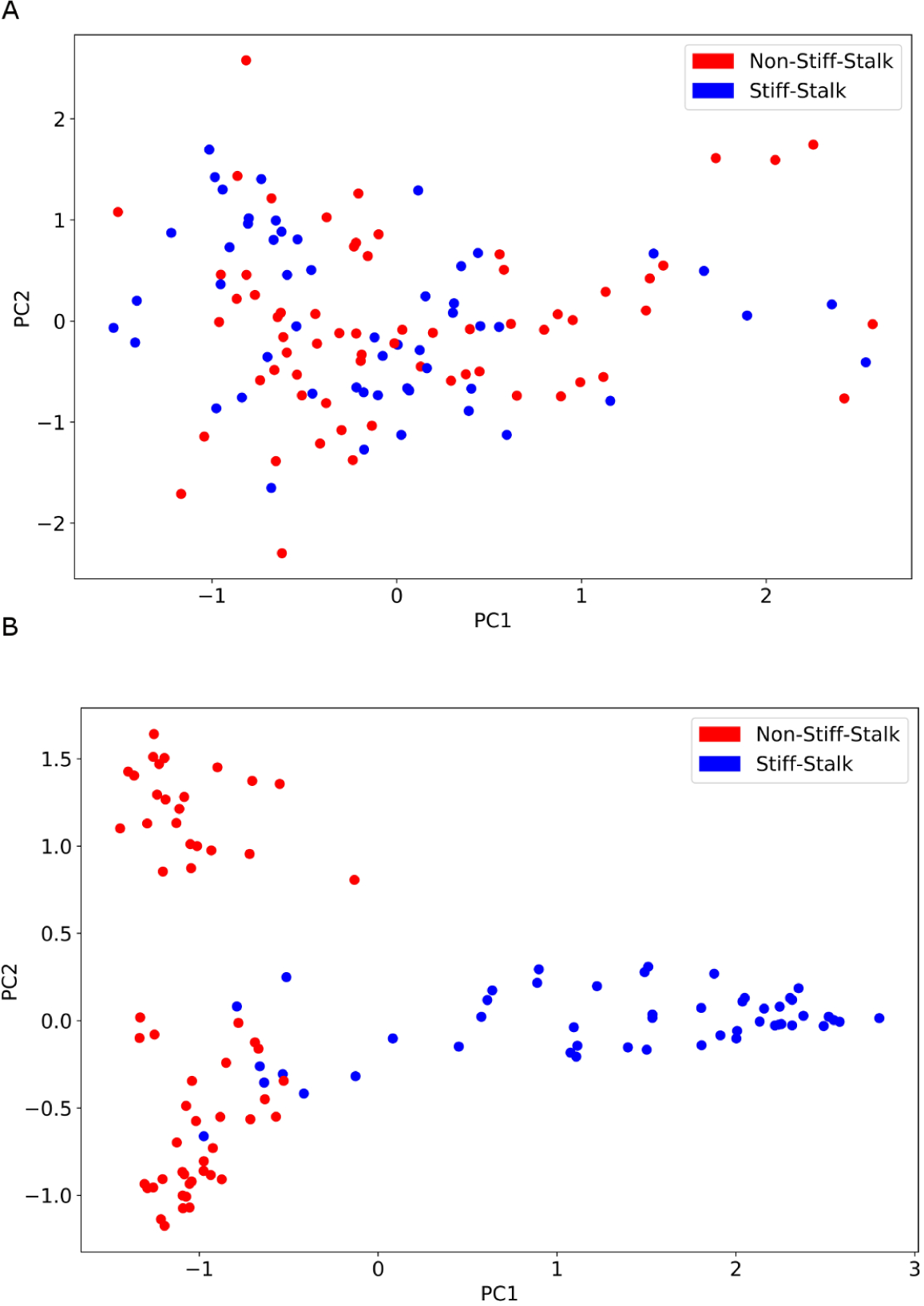
Population structure in the two heterotic groups of maize. In (A) the first two dimensions of a PCA analysis are used to describe the population structure as inferred by repeat content of the genome (i.e. abundance estimates of the clusters). In (B) PCA is used to infer population structure, but using genome wide SNP markers.

The population structure observed in maize ex-PVP breeding germplasm is expected and is reflective of the reciprocal recurrent selection breeding methods used to drive heterosis between heterotic pools [34]. This trend is not observed when global divergence in repeat content is used to establish clusters. This difference indicates that allele frequency alterations caused by selection or drift do not result in a corresponding change in global repeat abundance. For example, if, at a given site, segregating haplotypes contain similar repeat content but vary in SNP content, then selection for beneficial SNP alleles could change SNP frequencies within the population but have limited effect global repeat profiles. This possibility is supported by the observation that current elite maize germplasm pools were developed from a collection of similar landrace founders [34], leaving the possibility that repeat content was rather uniform across the founder population and has not yet had time to diverge in modern breeding populations.

This result is not supported by observation that inbred maize genomes show high variability in repeat co-linearity in and around genes [28] and across the genome [36]. This local repeat variation should track with linked SNPs and become evident in global repeat profiles. Furthermore, the difference in SNP and repeat abundance analyses could be driven by the ability of repeats, particularly transposable elements, to break linkage associations, limiting the effect of breeding history on repeat distribution and abundance. Lastly, the possibility remains that graph-based clustering and quantification is not adequate to detect the variation we know exists. As mentioned previously, this seems unlikely given the wealth of previous studies where variation in repeat content has been readily detected [9, 5, 11, 14], and indeed, can follow closely relationships between taxa [15].

## 4 Conclusion

We demonstrate the utility of graph-based clustering for repeat identification and quantification in maize, and develop a kmer-based approach to analyzing repeat content distribution across chromosomes *in silico*. Kmer painting of repeats reveals various chromosomal distribution patterns and we provide evidence that repeat clusters accumulate along chromosomes in a cluster dependent manner that can be negatively, or positively associated with local gene density. This novel analysis provides an additional avenue of investigation for researchers using graph-based repeat identification.

Additionally, we reveal that genetic population structure, as indicated by genotype data, distinguishes two well-known heterotic groups on elite maize, whereas global repeat populations are homogeneous between the two populations. These observations suggest that the highly dynamic repeat fraction of the inbred maize genome has not diverged in concert with SNP profiles in the two populations, SNP and repeat abundance profiles are uncoupled.

## Declarations

### 4.1 Ethics Approval and Consent to Participate

Not Applicable.

### 4.2 Consent for publication

Not Applicable.

### 4.3 Availability of Data and Material

The datasets generated and/or analysed during the current study are available in the NCBI SRA repository, https://submit.ncbi.nlm.nih.gov.

## Acknowledgements

We thank Justin Gerke, Eli Rodgers-Melnick, Steve McKay, Stephanie Coffman, Ming Yang and Joseph Evans for thoughtful discussion during the preparation of this manuscript.

## Additional Files

Additional file 1 — List of ex-PVP inbreds used in this study

A list of the names for ex-PVP inbreds used in this study.

Additional file 2 — Cluster annotation information

File listing clusters and their associated annotation as determined by sequence matches to RepBase

Additional file 3 — Cluster read counts by inbred

A file listing cluster name and the number of reads from each inbred that contributed to the given cluster.

Additional file 4 — Genome-wide kmer distribution in B73 v3.23

The genome-wide K-mer distribution for each of the 25 largest repeat clusters. A Kernel density estimation is indicated by color, and associated gene density estimates are also given.

Additional file 5 — Genome-wide kmer distribution in B73 v4.0

The genome-wide K-mer distribution for each of the 25 largest repeat clusters. A Kernel density estimation is indicated by color, and associated gene density estimates are also given.

Additional file 6 — Genome-wide kmer distribution in PH207

The genome-wide K-mer distribution for each of the 25 largest repeat clusters. A Kernel density estimation is indicated by color, and associated gene density estimates are also given.

